# How clustering promotes biodiversity: A solution to the plankton paradox

**DOI:** 10.1101/2023.12.17.572061

**Authors:** Karen K. Huruntz, Lilit E. Ghukasyan, Gor A. Gevorgyan

## Abstract

From a mathematical perspective, randomly assembled species-rich competitive communities exhibit a high degree of instability, as described by May’s stability theorem. This suggests that actual ecological communities should possess a significant level of organization in their competitive network structure, often with a modular topology, to withstand May’s instability. It is also known that a species-saturated ecological community driven by competitive Lotka–Volterra equations, following a path of competitive exclusion, lingers in a long-term state with species congregated in niche space into well-separated clusters, thereby competing in a modular manner. Without regard to origin, we characterize the modular network topology of interspecific competition in such a community as a hierarchical one, representing interspecific competition and the associated fragmentation of niche space among community species as a two-tiered process. That is, in pairs between species within individual groups (clusters) of ecologically related species, on the one hand, and collectively between these groups as holistic ecological units – ecological species, on the other hand. To illustrate this hierarchical organization, we introduce a generalized multi-resource multi-species competition model that recognizes this topology at the basis of its own network structure of interrelated governing equations of species dynamics. This kind of generalization substantially reduces the competitive exclusion constraints on species coexistence, offering promising potential for a conceptual resolution of the plankton paradox.

## INTRODUCTION

The plankton paradox, a term coined by Hutchinson (1961), refers to the enormous diversity of phytoplankton species observed virtually in all aquatic ecosystems. Meanwhile, a straightforward understanding of the principle of competitive exclusion (Hardin 1960; Armstrong and McGehee 1980), the core tenet of theoretical ecology, teaches that in competitive equilibrium a number of species competing for a fewer number of resources cannot coexist. Indeed, the aquatic environment is apparently too homogeneous to allow different species of phytoplankton, competing for light and, in order of magnitude, a dozen mineral resources, to be settled in different compartments in niche space, unless the principle “one niche – one species” is violated. One way out of this difficulty is hinted at in the very formulation of the principle of competitive exclusion, which oriented Hutchinson (1961) to seek a solution to the problem in that varying environmental conditions prevent the formation of competitive equilibrium in a phytoplankton community, thereby suspending the extrusion of less adapted species. Another reason for suspending competitive exclusion was hypothesized to be the internal chaotic dynamics in multispecies competitive communities (Huisman and Weissing 1999; Scheffer et al. 2003; Benincà 2008). Over the past six decades, other solutions to the problem and their variations have been discussed in the literature (Wilson 1990; Roy and Chattopadhyay 2007; Behrenfeld and Bisson 2024), with an equally remarkable diversity of proposed mechanisms. While most allow the coexistence of a few more competitors in addition to what is allowed by competitive exclusion, only a few highly idealized theories allow for the coexistence of hundreds of species limited by a small number of resources. In the absence of a universally agreed-upon resolution to the issue, we outline here another conceptual solution that, besides, fits well within the framework of the principle of competitive exclusion, rather than specifying mechanisms that mitigate competitive exclusion in species-rich communities. However, this comes at the expense of rethinking the principle of competitive exclusion to address the question regarding the specific meaning of the term “species” in this context. This issue holds relevance at least due to the multiplicity of species concepts (Aldhebiani 2018), such as biological, morphological, ecological, and phylogenetic concepts, among others. To our knowledge, this matter has never arisen before. However, once formulated, the answer suggests itself – ecological species, which we will promote below with ensuing consequences.

## CONCEPTUAL FRAMEWORK

At the first step of our reasoning, we recall the so-called May instability (Allesina and Tang 2015). Somewhat folkloric, somewhat under the authorship of Elton (1958), MacArthur (1955) and others, the dominant view was that the stability of an ecosystem positively correlates with its biodiversity and complexity. However, in the early 1970s, May (1972) sharply challenged this opinion, from a mathematical perspective showing exactly the opposite. May himself, referring to Gardner and Ashby’s (1970) computations, saw a solution to the problem his analysis initiated in the block structure of the community-matrix of an ecosystem (Grilli et al. 2016).

More recent numerical simulations of the dynamics of the species-rich Lotka-Volterra community with interspecific competition quantified by niche overlap have indeed shown a lumpy distribution of species along a niche axis (Scheffer and van Nes 2006; Fort et al. 2009; Pigolotti et al. 2009), which actually confirms May’s rather general surmise in a particular embodiment. We hypothesize that this is also the case with phytoplankton communities, all the more so because there are empirical evidences to support this (Vergnon et al. 2009). Indeed, drawing on the correlations between morphological properties and functional traits of phytoplankton species (Kruk and Segura 2012), a number of studies (Segura et al. 2011, 2013; Graco-Roza et al. 2021) have suggested that the multimodal pattern observed in species size distribution within phytoplankton communities reflects a clustered arrangement of species along niche gradients.

The following question guides our further reflection. Should we think of this phenomenon of grouping of species in niche space as something transient and purely quantitative along the path of competitive exclusion? With a positive answer, one may trace the dynamics of the community to the state of bankruptcy of individual clusters to single species (Scheffer and van Nes 2006). Not what we seek. Otherwise, in order to account for the suppression of competitive exclusion at this stage, mechanisms akin to those mentioned at the outset will have to be invoked again, albeit with a more favorable prospect. However, there is another option to untangle the dilemma outlined. Namely, we ought to take this state for granted, by and large, without regard to the mechanisms that brought the system to this state, whether they involve May’s instability (Grilli et al. 2016), competitive exclusion (Haraldsson and Thébault 2023), or evolutionary assembly of a community (Bonsall et al. 2004; Kashtan and Alon 2005), instead accepting the modular arrangement of species in niche space into clusters as a principal internal determinant of subsequent community dynamics.

The general scheme of mathematical models of the dynamics of ecological communities is such that each species of the community is assigned its own dynamic equation from a finite class of equations, say logistic ones, while the interaction between species is modeled by appropriately linking the governing equations corresponding to the interacting species. This way, the network topology of interspecific relationships is mirrored in the network of interrelations among the species governing equations, thereby imposing a predetermined topological constraint on the model. As a consequence, even if the niche structure of the model community shows a clear tendency towards a new topological order, the imposed dynamics by itself cannot provide this topological transition, but rather suggest the possibility of such a change. And seeing behind the dynamics of the community a transition to a qualitatively new topological state, not potential, but actual, we have to restructure the network of interrelations between the governing equations of species, conforming it with the newly emerging niche topology of the community. Below we are going to implement this program for communities with a modular network of interspecific competition.

The intensity of competition among organisms is contingent upon the resemblance of their requirements for shared resources of limited supply (Welden and Slauson 1986; Adler et al. 2018). The classification of competitive relationships as either intraspecific or interspecific reflects precisely the qualitative distinction that should exist in these requirements of individuals from the same versus different species. However, following this line of reasoning, we contend that it would also be warranted to make a qualitative distinction between competition among species within the same cluster and those from different clusters in competitive communities featuring a well-defined modular niche structure. Thus, when dealing with an ecological community comprising species clumpy distributed in niche space, and quantifying competition between species based on the overlap of their ecological niches, the degree to which interspecific competition contrasts within and between clusters largely depends on the degree of modularity of the community niche structure. In the asymptote of high modularity of the community’s competitive network, this contrast will obviously acquire a qualitative character, which makes it quite reasonable to accept the same idea even with a moderately high level of modularity of the competitive network.

However, the subclassification of interspecific competition as either intra-cluster or inter-cluster competition at this stage of the reasoning holds only nominal significance. To give it practical meaning, we will introduce the main assumption of the study. For this purpose, we first invoke the concept of ecological species (Andersson 1990), deeming it as the most appropriate notion for characterizing the species clusters we are talking about. Indeed, these clusters comprise species with closely adjacent and significantly overlapping realized niches, which, by the same token, are greatly narrowed due to intense competition within the clusters. All of this clearly suggests that the fundamental niches of species within individual clusters significantly coincide, thus fulfilling the criterion of an ecological species (Andersson 1990).

With the above considerations, we are now ready to introduce the main assumption of the study, designed to demonstrate our comprehension of the network topology of competitive relationships in competitive ecological communities with a clumpy arrangement of species in niche space. Namely, species within individual clusters compete in a general manner, while inter-cluster competition of species proceeds as competition between ecological species acting as holistic ecological units and represented by these very clusters. This assumption outlines a certain hierarchical topology for modular competitive networks (Fig. 1).

**Figure 1.**
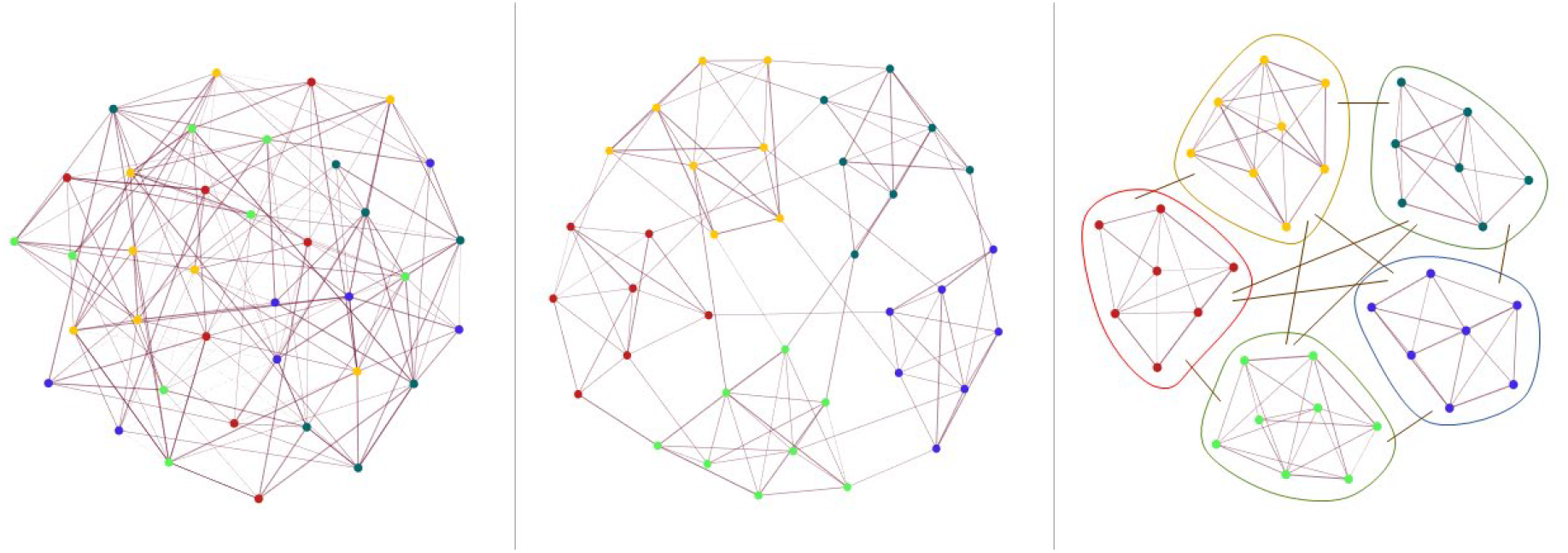
**Network structure of a community** with a general (random) distribution of competitive relationships between species (left panel); with a clear cluster aggregation of species, when species compete more intensely within clusters and less intensely between clusters (middle panel); with a hierarchical topology of competitive links, when species compete in a general manner in clusters, while inter-cluster species competition proceeds as inter-cluster competition with species being amalgamated in holistic groups – ecological species (right panel).

It is important to acknowledge that this assumption is somewhat conceptual in nature, and as such, its validation should be based on empirical evaluation of the ensuing consequences, rather than on logical reasoning derived from established frameworks. However, some rationale may be inferred from general considerations. When examining exploitation competition between different species, it is the species themselves or their populations that serve as the fundamental units in these competitive relationships. And if the concept of species in this particular context is interpreted solely as a means to identify these units of competitive relationships, it becomes entirely justifiable to disregard any species criteria that are not directly related to the species’ abilities to exploit the contested shared resources. Instead, the focus should be on the species criteria that are relevant from the contextual perspective, namely those that identify similarities and differences among groups of individuals in terms of resource utilization. This is precisely what may justify the recognition of ecological species as the primary unit of interspecific competition. However, such reasoning delineates only the upper coarse-grained level of competition in an approximation that disregards fine-grained ecological differences between biological species within individual ecological niches of ecological species. Further recognition of this fact distinguishes another level of interspecific competition, this time between biological species, that, however, does not extend to the entire community, but is localized within the niches of individual clusters of ecologically related species.

Overall, the following picture emerges. In the case under consideration, interspecific competition and the resulting fragmentation of niche space have to be thought of as a sequential hierarchical process, not in the course of time, but along the path of reasoning. The coarsegrained level of fragmentation demarcates compartments in niche space for the quartering of clusters, comprising ecologically related biological species whose ecological traits are indistinguishable at this level of detail between species in the same cluster and clearly distinguishable between species from different clusters, and competing as integral groups – ecological species – with other peer groups for a vital amount of niche space. Further finegrained fragmentation, in turn, occurs in individual niche compartments of ecological species as a result of competition between biological species that make up a given ecological species. It is also worth noting the somewhat self-similar nature of the above hierarchical representation of modular competitive networks, when the lower level of a community competitive network repeats, in a sense, the upper level, while also providing the upper-level nodes with an internal network structure. Should this replication continue indefinitely, a fractal structure could emerge. However, the prospect of its application in a multi-level format for ecological intentions is scarcely warranted.

## MATHEMATICAL CLARIFICATION

To further clarify the above from a mathematical perspective, we define a hierarchical extension of the Lotka-Volterra type multi-resource competition model to explore the constraints of competitive exclusion on the maximum number of coexisting species in a hierarchically structured competitive community. In general, the Lotka-Volterra equations, along with their nonlinear and stochastic extensions and variations, are widely regarded as a standard framework for examining the theoretical foundations of competitive exclusion (Chesson 2000, 2018). Moreover, it was Volterra (1926) who first explicitly articulated the competitive exclusion principle based on his theoretical disquisitions, antedating Gause’s (1934) model experiments. However, early reflections on competitive exclusion primarily concentrated on species vying for a single resource. Extension to the case of multi-resource competition was achieved in the 1960s in a number of studies (MacArthur and Levins 1964; Rescigno and Richardson 1965; Levins 1968; Levin 1970) based on slightly different premises (Armstrong and McGehee 1980). The common quantitative conclusion of these studies, which remains pertinent up to now, is that at steady state the number of competing species cannot exceed the number of limiting factors, shared resources or niches (Armstrong and McGehee 1980). This is what gives a quantitative dimension to the problem of species coexistence, in particular the plankton paradox. In what follows, we intend to translate this result into the context of the hierarchical organization of a competitive network described above. We will build on the model and its analysis presented in the classic paper by Levin (1970), which has significantly shaped our current comprehension of competitive exclusion in multi-resource multi-species communities. However, this choice is largely arbitrary, as will be discussed towards the end.

First, we define the effective abundance of the *i*th cluster (*M*_*i*_) as the sum of the abundances of all species in this cluster (*N*_*i,j*_): *M*_*i*_ = Σ_*j*_ *α*_*i,j*_*N*_*i,j*_, weighted by coefficients reflecting the relative per capita capacities of species to exploit shared resources. The fact is that clusters unite species that share similar ecological traits at the population level, such as having equal proportions in their needs for various common resources and possessing similar capacities to exploit these resources. However, this does not ensure uniformity in ecological traits per capita. In such an instance, we have to assign equal values to all coefficients.

By employing Levin’s (1970) framework, we now model the coarse-grained level of competition between clusters through the following system of equations:

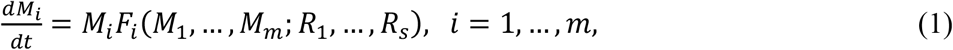

where *m* denotes the number of clusters in the community, and the variables (parameters) *R*_1_, …, *R*_*s*_ bring into play the overall set of resources (factors) shared at the level of inter-cluster competition. The system (1) governs the dynamics of the community at the inter-cluster level. It determines the course of competition and the outcome of competitive displacement of less adapted clusters competing as holistic ecological units for coarse-grained fragmentation of niche space. This ultimately determines the number of clusters that persist in the competitive equilibrium and their effective abundances. Within these effective abundances of clusters emanating from system (1), further fine-grained competition occurs among the species of each individual cluster for the same set *R*_1_, …, *R*_*s*_ of resources; although, in general, the actual subset of shared resources may vary from cluster to cluster and at the level of inter-cluster competition. Referring again to Levin’s (1970) model and fixing the cluster index *i*, this competition is modeled by the following system of equations:

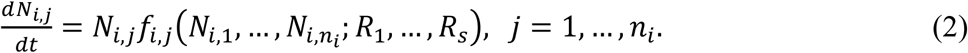

where *n*_*i*_ represents the number of species in the *ii*th cluster.

With the provisos above, the model (1) and (2) constitutes a system of interrelated differential equations with a hierarchical topology of the network of connections between them, which is what fundamentally distinguishes it from the archetypal one. The system (1) provides a description of how species from different clusters interact collectively, whereas equations (2) with a fixed cluster index *i* only account for the interaction between species within that particular cluster. In this regard, it is worth noting that equations (2) are independent of each other for different values of the cluster index *i*, except for their connection through system (1). The system (1), in turn, is related to equations (2) only through the definition of the effective abundances of clusters, connecting them with the abundances of species within individual clusters.

Refraining from delving into the mathematical analysis of the model defined by equations (1) and (2), we only mention one feature of it, which is directly related to the plankton paradox and the principle of competitive exclusion and can be deduced by analogy with the long-known properties of the modified Lotka–Volterra multi-resource competition model (Levin 1970; Armstrong and McGehee 1980). Thus, the system of equations (1), as well as each subsystem of the system (2) obtained by fixing the cluster index *i*, from a mathematical point of view repeats that system, as it has been designed consciously. Therefore, abiding by the analysis of Levin (1970), we have to move from shared resources to limiting factors as a minimal complete set of independent arguments for the functions *F*_*i*_ and *f*_*i,j*_, with no smaller collection of variables being sufficient to serve this purpose:

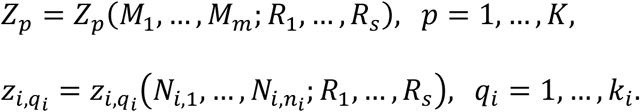

With these variables *Z*_*p*_, representing *K* limiting factors at the level of inter-cluster competition, and 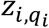, representing *k*_*i*_ limiting factors for interspecific competition within the *i*th cluster, the system (1) and (2) takes the following form:

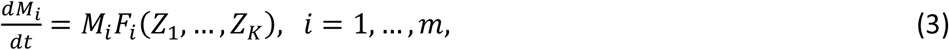

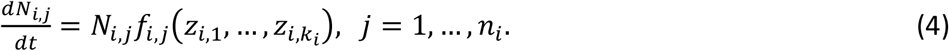

Dating back to Volterra’s (1926) seminal work, it has been a common practice to treat functions *F*_*i*_ and *f*_*i,j*_ as linear functions of their arguments. Assuming this premise and further replicating Levin’s (1970) analysis separately for the system (3), we will inevitably reach the same conclusion, which can be paraphrased in a common manner as follows: the state of competitive equilibrium of the community, cyclic or static, can accommodate no greater number of clusters of species than the number of limiting factors at this level: *m* ≤ *K*. As for the system of equations (4), for a fixed cluster index *i*, it controls the interconnected dynamics of species within this particular cluster, but with one additional condition linearly relating the abundances of species in this cluster to the effective size of the cluster emanating from (3). However, it is quite obvious that this extra clause cannot in any way mitigate the just-mentioned constraint on the maximum number of species in equilibrium implied by the system (4) without this extra clause: *n*_*i*_ ≤ *k*_*i*_, but perhaps tighten it. Overall, we conclude that the equilibrium of the model under consideration can harbor in total at most as many species as 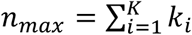.In the simplest situation, when the number of limiting factors is the same at the level of inter-cluster competition and in each cluster, the competitive exclusion constraint boils down to *n* ≤ *k*^2^. This stands in stark contrast to the competitive exclusion constraint on the maximum number of community species established for flat (non-hierarchical) competitive systems as *n* ≤ *k*. This expresses the hierarchical structure of competitive relations within the proposed model. Given a dozen limiting factors and hundreds of phytoplankton species, this is quite enough to conceptually resolve the plankton paradox.

The above consideration does not address the question of the achievability of the limit imposed on the maximum number of species in an equilibrium community. However, it should be noted that this issue has been largely silent in other theoretical studies of the principle of competitive exclusion (MacArthur and Levins 1964; Rescigno and Richardson 1965; Levins 1968; Levin 1970; Armstrong and McGehee 1980) and seems to be a relevant open question.

## CONCLUDING REMARKS

In conclusion, we would like to emphasize that the main thesis of the above concerns not the equations with which we have embodied the main message of this work, but the hierarchical network structure of competitive connections that we attribute in particular to phytoplankton communities. Moreover, we could base our hierarchical structure on another model of competitive exclusion, such as that by Rescigno and Richardson (1965) or Levins (1968). Provided that this model upholds a constraint *n* < *k* for a flat competitive system, we would arrive at the same result regarding the constraint of competitive exclusion on the maximum number of community species, expressed as *n* < *k*^2^. Furthermore, we have the option to forgo utilizing a specific underlying model altogether and instead adopt the *n* < *k* constraint across the various levels of hierarchical competition outlined in the study. This will again result in the same outcome.

In the context of the plankton paradox itself, we also note that the above resolution does not in any way contradict most previously studied approaches to solving this problem, such as those associated with intrinsic dynamic chaos in phytoplankton communities (Huisman and Weissing 1999; Benincà 2008) or variable external conditions (Hutchinson 1961), but rather can incorporate them into itself with great harmony, leading to a more sustainable solution to the problem.

## ACKNOWLEDGMENTS

This work was partly supported by the Science Committee of MESCS RA in the frame of the research project no. 21T-1F222.

## REFERENCES

Adler, P. B., Smull, D., Beard, K. H., Choi, R. T., Furniss, T., Kulmatiski, A., Meiners, J. M., Tredennick, A. T., Veblen, K. E. (2018). Competition and coexistence in plant communities: intraspecific competition is stronger than interspecific competition. Ecology Letters 21, 1319–1329. 10.1111/ele.13098

Aldhebiani A. Y. (2018). Species concept and speciation. Saudi Journal of Biological Sciences 25, 437–440. 10.1016/j.sjbs.2017.04.013

Allesina, S., Tang, S. (2015). The stability–complexity relationship at age 40: a random matrix perspective. Population Ecology 57, 63–75. 10.1007/s10144-014-0471-0

Andersson, L. (1990). The driving force: species concepts and ecology. Taxon 39, 375–382. 10.2307/1223084

Armstrong, R. A., McGehee, R. (1980). Competitive exclusion. The American Naturalist 115, 151–170. 10.1086/283553

Behrenfeld, M. J., Bisson, K. M. (2024). Neutral theory and plankton biodiversity. Annual Review of Marine Science 16, 283–305. 10.1146/annurev-marine-112122-105229

Benincà, E., Heerkloss, R., Jöhnk, K. D., Branco, P., van Nes, E. H., Scheffer, M., Ellner, S. P. (2008). Chaos in a long-term experiment with a plankton community. Nature 451, 822–825. 10.1038/nature06512

Bonsall, M. B., Jansen, V. A. A, Hassell, M. P. (2004). Life history trade-offs assemble ecological guilds. Science 306, 111–114. 10.1126/science.1100680

Chesson, P. (2000). Mechanisms of Maintenance of Species Diversity. Annual Review of Ecology, Evolution, and Systematics 31, 343–366. 10.1146/annurev.ecolsys.31.1.343

Chesson, P. (2018). Updates on mechanisms of maintenance of species diversity. Journal of Ecology 106, 1773–1794. 10.1111/1365-2745.13035

Elton, C. S. (1958). The Ecology of Invasions by Animals and Plants. London: Methuen. 10.1007/978-1-4899-7214-9

Fort, H., Scheffer, M., van Nes, E. H. (2009). The paradox of the clumps mathematically explained. Theoretical Ecology 2, 171–176. 10.1007/s12080-009-0040-x

Gardner, M. R., Ashby, W. R. (1970). Connectance of large dynamic (cybernetic) systems: critical values for stability. Nature 228, 784. 10.1038/228784a0

Gause, G. F. (1934). The struggle for existence. Baltimore: Williams & Wilkins.

Graco-Roza, C., Segura, A. M., Kruk, C., Domingos, P., Soininen, J., Marinho, M. M. (2021). Clumpy coexistence in phytoplankton: the role of functional similarity in community assembly. Oikos 130, 1583–1597. 10.1111/oik.08677

Grilli, J., Rogers, T., Allesina, S. (2016). Modularity and stability in ecological communities. Nature Communications 7, 12031. 10.1038/ncomms12031

Haraldsson, M., Thébault, E. (2023). Emerging niche clustering results from both competition and predation. Ecology Letters 26, 1200–1211. 10.1111/ele.14230

Hardin, G. (1960). The competitive exclusion principle. Science 131, 1292–1297. 10.1126/science.131.3409.1292

Huisman, J., Weissing, F. J. (1999). Biodiversity of plankton by species oscillations and chaos. Nature 402, 407–410. 10.1038/46540

Hutchinson, G. E. (1961). The paradox of the plankton. The American Naturalist 95, 137– 145. 10.1086/282171

Kashtan, N., Alon, U. (2005). Spontaneous evolution of modularity and network motifs. Proceedings of the National Academy of Sciences 102, 13773–13778. 10.1073/pnas.0503610102

Kruk, C., Segura, A. M. (2012). The habitat template of phytoplankton morphology-based functional groups. Hydrobiologia 698, 191–202. 10.1007/s10750-012-1072-6

Levin, S. A. (1970). Community equilibria and stability, and an extension of the competitive exclusion principle. The American Naturalist 104, 413–423. 10.1086/282676

Levins, R. (1968). Evolution in changing environments. Some theoretical explorations. Princeton: Princeton University Press.

MacArthur, R. H. (1955). Fluctuations of animal populations and a measure of community stability. Ecology 36, 533–536. 10.2307/1929601

MacArthur, R., Levins, R. (1964). Competition, habitat selection, and character displacement in a patchy environment. Proceedings of the National Academy of Sciences 51, 1207– 1210. 10.1073/pnas.51.6.1207

May, R. M. (1972). Will a large complex system be stable? Nature 238, 413–414. 10.1038/238413a0

Pigolotti, S., López, C., Hernández-García, E., Andersen K. H. (2009). How Gaussian competition leads to lumpy or uniform species distributions. Theoretical Ecology 3, 89– 96. 10.1007/s12080-009-0056-2

Rescigno, A., Richardson, I. W. (1965). On the competitive exclusion principle. The Bulletin of Mathematical Biophysics 27, 85–89. 10.1007/BF02477264

Roy, S., Chattopadhyay, J. (2007). Towards a resolution of ‘the paradox of the plankton’: A brief overview of the proposed mechanisms. Ecological Complexity 4, 26–33. 10.1016/j.ecocom.2007.02.016

Scheffer, M., Rinaldi, S., Huisman, J., Weissing, F. J. (2003). Why plankton communities have no equilibrium: solutions to the paradox. Hydrobiologia 491, 9–18. 10.1023/A:1024404804748

Scheffer, M., van Nes, E. H. (2006). Self-organized similarity, the evolutionary emergence of groups of similar species. Proceedings of the National Academy of Sciences 103, 6230– 6235. 10.1073/pnas.0508024103

Segura, A. M., Calliari, D., Kruk, C., Conde, D., Bonilla, S., Fort, H. (2011). Emergent neutrality drives phytoplankton species coexistence. Proceedings of the Royal Society of London Series B: Biological Sciences 278, 2355–2361. 10.1098/rspb.2010.2464

Segura, A. M., Kruk, C., Calliari, D., García-Rodriguez, F., Conde, D., Widdicombe, C. E., Fort, H. (2013). Competition drives clumpy species coexistence in estuarine phytoplankton. Scientific Reports 3, 1037. 10.1038/srep01037

Vergnon, R., Dulvy, N. K., Freckleton R. P. (2009). Niches versus neutrality: uncovering the drivers of diversity in a species-rich community. Ecology Letters 12, 1079–1090. 10.1111/j.1461-0248.2009.01364.x

Volterra, V. (1926). Variazioni e fluttuazioni del numero d’individui in specie animali conviventi. Memoria della Reale Accademia Nazionale dei Lincei 2, 31–113.

Welden, C. W., Slauson, W. L. (1986) The intensity of competition versus its importance: an overlooked distinction and some implications. The Quarterly Review of Biology 61, 23– 44. 10.1086/414724

Wilson, J. B. (1990). Mechanisms of species co-existence: twelve explanations for Hutchinson’s ‘Paradox of the Plankton’: evidence from New Zealand plant communities. New Zealand Journal of Ecology 13, 17–42.

